# Remarkable genetic diversity of *Trypanosoma cruzi* and *Trypanosoma rangeli* in two localities of southern Ecuador identified via deep sequencing of mini-exon gene amplicons

**DOI:** 10.1101/542993

**Authors:** Jalil Maiguashca Sánchez, Salem Oduro Beffi Sueto, Philipp Schwabl, Mario J. Grijalva, Martin S. Llewellyn, Jaime A. Costales

**Affiliations:** Center for Research on Health in Latin America, School of Biological Sciences, Pontifical Catholic University of Ecuador, Quito, Ecuador; Institute of Biodiversity, Animal Health and Comparative Medicine, University of Glasgow, Glasgow G128QQ, UK; Infectious and Tropical Disease Institute, Department of Biomedical Sciences, Heritage College of Osteopathic Medicine, Ohio University, Athens, OH, USA

## Abstract

*Trypanosoma cruzi*, causative agent of Chagas disease, *and T. rangeli* are kinetoplastid parasites endemic to Latin America. Although closely-related to *T. cruzi* and capable of infecting humans, *T. rangeli* is non-pathogenic. Both parasite species are transmitted by triatomine bugs, and the presence of *T.* rangeli constitutes a confounding factor in the study of Chagas prevalence and transmission dynamics. *T. cruzi* possesses high molecular heterogeneity: six discrete typing units (DTUs) are currently recognized. In Ecuador, TcI predominates while other DTUs are seldom reported. Here, infection by *T. cruzi* and/or *T. rangeli* in triatomine bugs from two communities of southern Ecuador was evaluated via PCR product size polymorphism of kinetoplast-minicircle sequences and the non-transcribed spacer region of the mini-exon gene. Additionally, mini-exon amplicons were deep-sequenced to analyze single nucleotide polymorphisms and mixed infections. As a control, mini-exon products from several monoclonal reference strains were included. *T. cruzi* genetic diversity was significantly greater in adult vectors than in nymphal stage V vectors. Among infected triatomines, deep sequencing revealed one *T. rangeli* infection (3%), 9 *T. cruzi* infections (27.3%) and 23 *T. cruzi* / *T. rangeli* mixed infections (69.7%), suggesting that *T. rangeli* prevalence has been largely underestimated in the region. Furthermore, deep-sequencing detected TcIV sequences in six samples (first TcIV record in southern Ecuador). Our data indicate that amplicon size analysis alone is not reliable for parasite identification/typing in mixed infections containing both *T. cruzi* and *T. rangeli*, or when multiple *T. cruzi* DTUs are present. Additionally, our analysis showed extensive overlap among the parasite populations present in the two studied localities (ca. 28 km apart); suggesting active parasite dispersal over the study area. Our results highlight the value of amplicon sequencing methodologies to clarify the population dynamics of kinetoplastid parasites in endemic regions and inform control campaigns in southern Ecuador.

**Author Summary:** Chagas disease, caused by *Trypanosoma cruzi*, is a vector-borne parasitosis affecting ~6 million people in Latin America. We studied the molecular diversity of *T. cruzi* and *T. rangeli*, a closely related yet harmless parasite, in two localities in southern Ecuador. Due to their similarity, *T. rangeli* constitutes a confounding factor when studying Chagas prevalence and transmission dynamics. We compared three different molecular techniques, two based on polymerase chain reaction and another including an additional deep-sequencing strategy to study intestinal content samples from infected triatomines (blood-feeding insects capable of harboring both parasites and transmitting the infection to mammalian hosts, including humans). Almost 70% of the samples were found to be co-infected with both parasite species, suggesting *T. rangeli*’s prevalence has been previously underestimated in southern Ecuador. Additionally, we report the presence of a *T. cruzi* variant known as TcIV for first time in southern Ecuador. No significant genetic differences were identified among parasite populations in the studied localities. Finally, higher *T. cruzi* diversity was found in adult triatomines than in earlier developmental stages. Our results highlight the value of deep-sequencing methodologies to clarify the population dynamics of kinetoplastid parasites in endemic regions and provide valuable information to support control campaigns.

## Introduction

Chagas disease (CD) is caused by the kinetoplastid protozoan parasite *Trypanosoma cruzi* and affects 6 to 7 million people worldwide [1]. This neglected disease is endemic of Latin America, where it poses risk of infection for 25 million people and kills an estimate 10,000 every year [2]. Transmission in endemic countries primarily occurs via contact with feces of an infected triatomine bug. However, other secondary mechanisms of transmission, such as blood transfusion, organ transplants, congenital transmission, consumption of infected food and laboratory accidents exist, and can cause infections in non-endemic regions where vectors are not present [3–5].

*Trypanosoma rangeli*, a related kinetoplastid parasite, has overlapping geographical distribution with *T. cruzi* and may also be found in the same triatomine vectors and mammalian hosts; mixed infections with both species have been reported [6–8]. Despite being considered nonpathogenic for humans, *T. rangeli* has epidemiological relevance due to its morphological and genetic similarity with *T. cruzi*, which may lead to false-positive results in microscopic and serologic tests used for diagnosis, the latter due to cross reactivity caused by shared antigenic determinants [9, 10]. Triatomines are also vectors for *T. rangeli*, although transmission occurs via the salivary route [11]. In Ecuador, little is known about this parasite. Employing PCR minicircle amplification to analyze over 3600 samples, ~10% of triatomines and mammals in Manabí and Loja provinces (central Ecuadorian coast and southern Ecuador, respectively) were reported to be infected with *T. rangeli*, while 1.25% presented *T. cruzi/T. rangeli* mixed-infections [12].

Invertebrate vectors for both parasites belong to the Triatominae subfamily, which includes 150 extant species, although a reduced number are capable of *Trypanosoma* transmission [13]. Sixteen species of triatomines are found in Ecuador [14, 15]. Among them, *Triatoma dimidiata, Triatoma carrioni, Panstrongylus chinai, Panstrongylus rufotuberculatus* and *Rhodnius ecuadoriensis* are epidemiologically relevant [8, 16]. *T. dimidiata* and *R. ecuadoriensis* have been the target of previous research and much more is known for these two species than any other triatomine in Ecuador [17–20]. *R. ecuadoriensis* shows a wide distribution that includes the western lowlands and the southern provinces of Ecuador, as well as northern Perú [21], both in domestic and peridomestic environments, where contact with humans also occurs [17, 22].

Infection by *T. cruzi* and/or *T. rangeli* in both vertebrates and invertebrates is frequently multiclonal, involving several parasite genotypes with genetically dissimilar profiles [23–26]. Multiclonal infections have been suggested to impact host immunity [27], drug resistance evaluation [28], transmission rate and population structure [29]. It has been proposed that parasite diversity accumulates over the lifespan of hosts, as a result of increased time of exposure to infection [29, 30]. In agreement with this hypothesis, Dumonteil and colleagues reported a positive correlation between *T. cruzi* diversity and the estimated number of infection events in *T. dimidiata* [31]. On the other hand, previous analysis of human infections in endemic areas showed no correlation between parasite diversity and patient age [29, 30].

During the last four decades, molecular patterns grouping *T. cruzi* strains and isolates in different clusters have been identified. Initially, three “zymodemes” were identified via multilocus enzyme electrophoresis [32–35]. With the advent of molecular techniques, a variety of molecular markers support the existence of six lineages or Discrete Typing Units (DTUs), commonly referred to as TcI - TcVI [36–40]. In Ecuador, TcI predominates in the central coast [19, 41] and southern highlands (Loja Province) [42]. Only two previous reports suggest the presence of parasites belonging to DTUs other than TcI in triatomines and chagasic patients [43, 44].

In the genus *Trypanosoma*, mini-exon genes are present in several tandemly arranged copies and encode the splice leader, a 35-nucleotide sequence translocated to the 5’-end of every newly synthesized mRNA [45, 46], in a process referred to as discontinuous transcription [47]. *T. cruzi* mini-exon genes include conserved, semi-conserved and highly variable regions, which have been used for phylogenetic analysis [48], discrimination between DTUs [49], population genetic inference [50], and more recently in diversity analysis within naturally infected mammalian hosts [51]. Nonetheless, it is not clear how well mini-exon sequence diversity and DTU identity are matched, and PCR-based assays targeting the mini-exon locus are not considered reliable for *T. cruzi* DTU detection and identification [51]. However, given its variability, sequencing of the intergenic spacer has been proven useful in the examination of population dynamics within a single DTU [52–56].

The advent of Next Generation Sequencing (NGS) represents a major innovation in biological and medical research. The last decade saw an explosion of NGS techniques and, by 2010, the capacity of data generation increased by several orders of magnitude compared to Sanger-sequencing technologies [57]. Amplicon sequencing, which involves the sequencing of the products of a targeted PCR has been widely adopted as a tool to explore microbial community diversity [58]. In the case of *T. cruzi*, the technique has helped to provide insights into multiclonal infections in humans [30] and has recently been used to assess relationships between triatomine gut microbiota parasite diversity, and vertebrate feeding sources of the insect [31].

In the present study, we aimed to evaluate the molecular diversity of *T. cruzi* and *T. rangeli* in two localities of southern Ecuador via analysis of the mini-exon gene. Alongside forty-six samples from Loja Province, 9 cloned isolates of reference strain DTUs I, III, IV, V and VI, were also analyzed in order to establish a correspondence between mini-exon diversity and DTU identity. Amplicon sequence reads generated by NGS were sorted into haplotype clusters (a set of sequences with 97% homology) to explore variability among intestinal content DNA, including measurement of genetic richness, diversity and possible divergence between localities in the study region. Moreover, the data set included samples isolated from different developmental stages of the vector to evaluate a potential correlation with parasite genetic diversity. We were also able to compare genotyping results from NGS and PCR product size polymorphism analysis in terms of sensitivity for discrimination between *T. cruzi* and *T. rangeli*.

## Materials and methods

### Sample panel and PCR typing

Archival DNA samples collected between years 2009-2013 by the Center for Research on Health in Latin America (CISeAL) were used. Details of entomological searches, intestinal content DNA extractions and corresponding approved protocols and collection permits have been reported elsewhere [7, 16]. Samples selected for the study had been previously genotyped by kinetoplast minicircle amplification using primer pairs S35 - S36 [59] and 121 - 122 [60], at which time they were deemed to be infected exclusively with *T. cruzi* (data not published). Fig 1 shows a representative gel displaying the banding patterns associated with *T. cruzi* and *T. rangeli* in this analysis.

**Fig 1.**
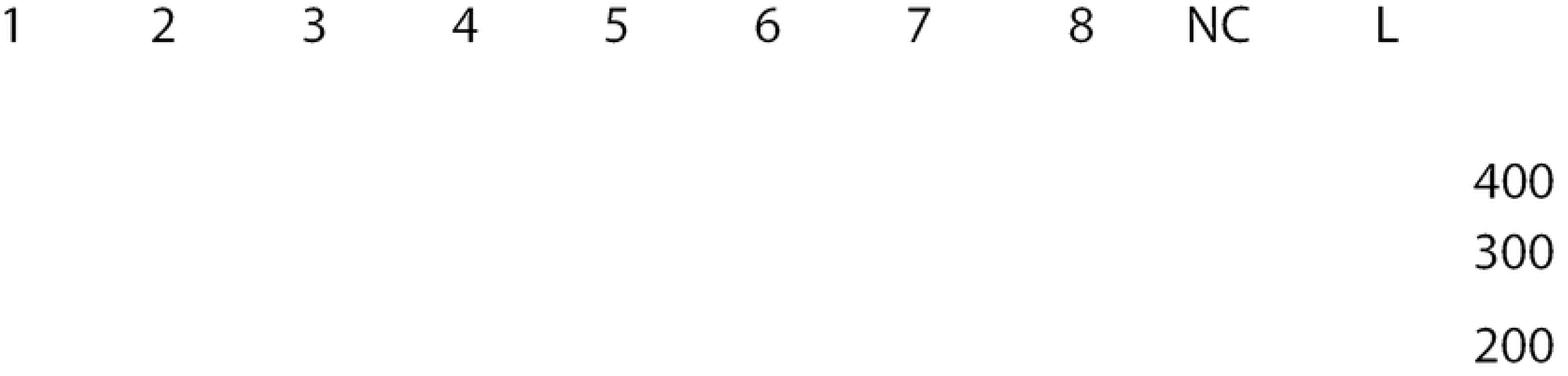
Banding patterns associated with *T. cruzi* and *T. rangeli* in the minicircle kinetoplast PCR analysis. Amplifications were performed using primers 121/122 [60] and electrophoretically separated in 2% agarose gels. Samples **1**: TCQ3084, *T. rangeli*; **2**: TCQ3087, *T. cruzi*; **3**: TBM3131*, *T. cruzi*; **4**: TBM3297, *T. cruzi*; **5**: TAZ3968, *T. cruzi*; **6**: THY4020, *T. rangeli*; **7**: THY3978, *T. cruzi*; **8**: THY4329, *T. cruzi*; **NC**: Negative control (ddH_2_0 as template); **L**: molecular weight ladder. *Only sample included in the subsequent analysis; banding pattern consistent with infection exclusively by *T. cruzi*.

Forty-six samples from two rural communities from southern Ecuador (Loja Province) were included in the sample panel: 31 from Bramaderos (− 4.0797, −79.8244) and 15 from Bellamaría (−4.2115, −79.6063) (Fig 2). To characterize the molecular diversity/multiclonality of the mini-exon gene of the parasites across vector developmental stages, we included samples obtained from 19 adult, 13 instar V, 11 instar III and 3 instar IV triatomine bugs.

**Fig 2.**
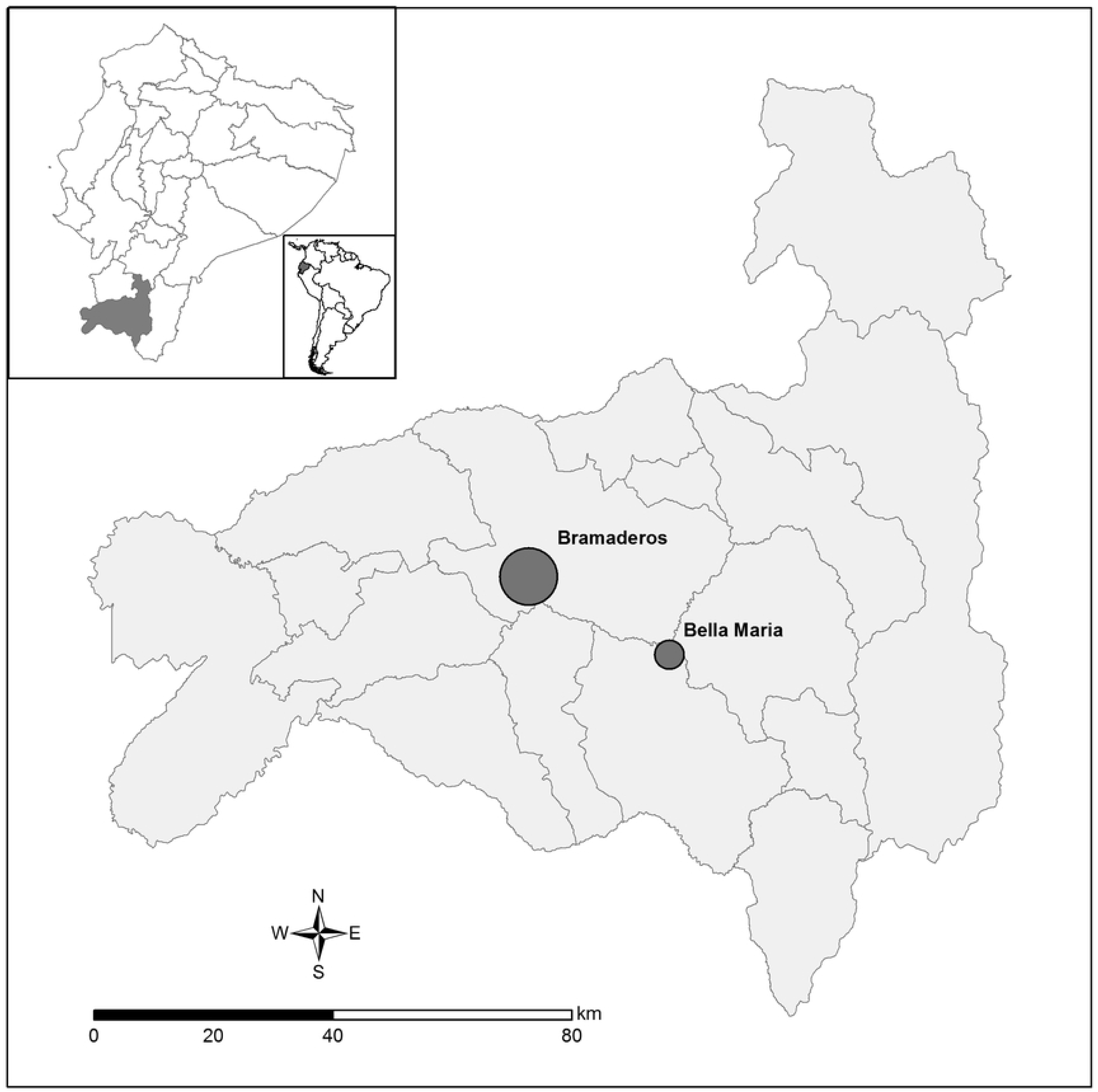
Study area. The two studied communities within Loja province in southern Ecuador are depicted. They are separated by approximately 28.3km. Circle diameter is proportional to sample size in each locality. Map generated with ArcGIS software, version 10.5, based on data freely available from the Military Geographic Institute of Ecuador (IGM, http://www.geoportaligm.gob.ec/portal/). Community location was obtained via GPS measurement.

The mini-exon gene is a multicopy target. Therefore, DNA from nine biologically cloned reference strains was included in the analysis in order to evaluate whether intra-clonal sequence diversity generated from the amplicons via NGS recapitulated the DTU to which the clones had been previously been assigned via classical methods [51].

### Multiplex PCR

PCR amplification of the non-transcribed spacer region of the mini-exon gene was performed employing five primers in a multiplex reaction to discriminate between zymodemes I, II, III and *T. rangeli*, as previously described [49]. In the current nomenclature zymodeme I is equivalent to DTU TcI, zymodeme II comprises DTUs II, V and VI, while zymodeme III corresponds to DTUs III and IV [40]. The amplified products were electrophoresed in 2% agarose gels and visualized with SYBR green under UV light. Samples coinfected with *T. cruzi* and *T. rangeli* displayed a two-band pattern (~210/~150bp) (Fig 3, M). In such cases, gel excision and purification were performed for each fragment. Subsequently, amplicons were sequenced by NGS technology as detailed below.

**Fig 3.**
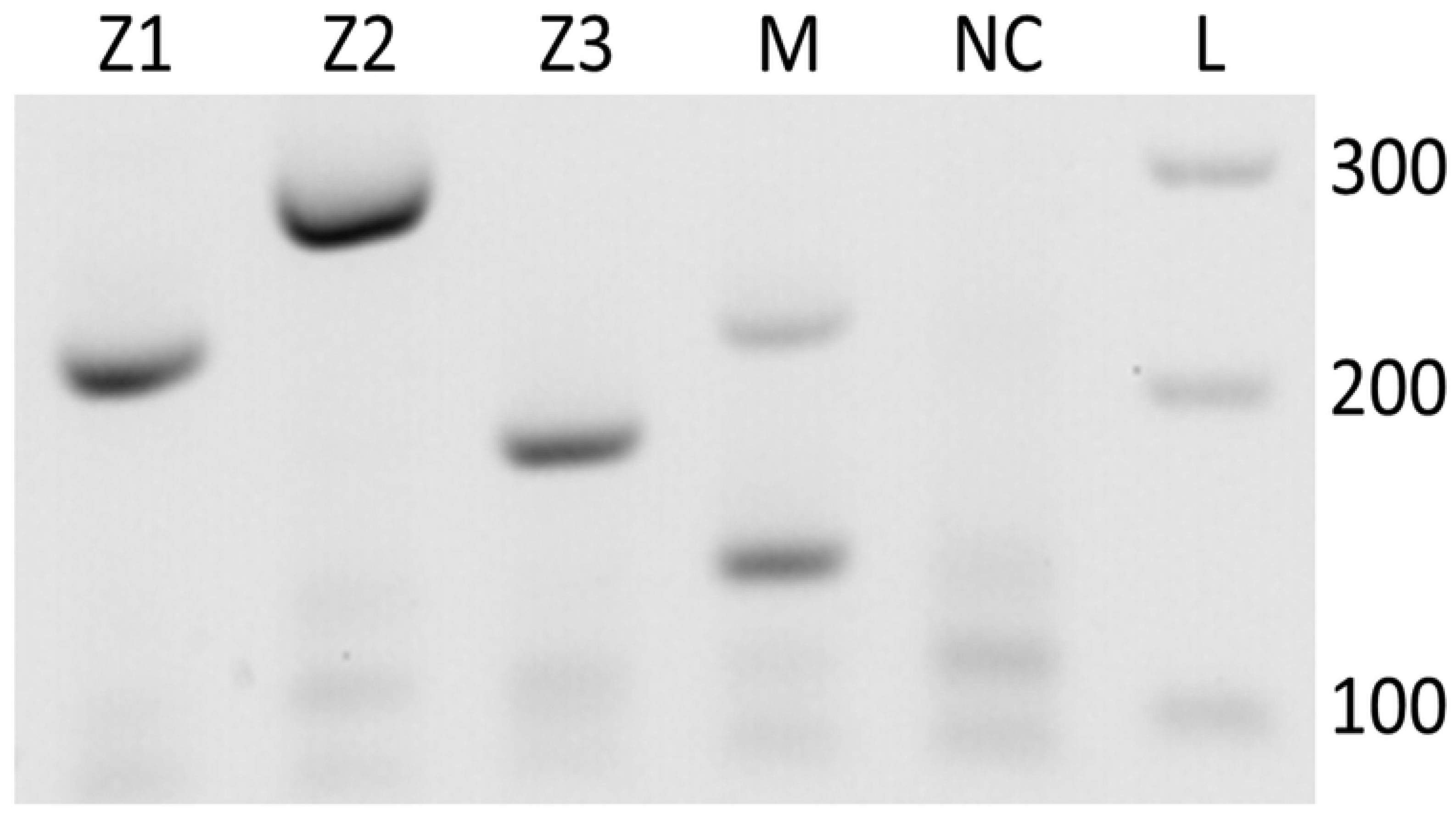
Representative PCR amplification pattern for the non-transcribed spacer of the mini-exon gene. PCR was performed as in [49], and amplicons were electrophoretically separated in 2% agarose gels. Banding patterns correspond to: **Z1**: *T.cruzi* zymodeme 1, equivalent to DTU TcI; sample TBR1391. **Z2**: *T. cruzi* zymodeme 2, corresponding to DTUs TcII, TcV and TcVI; Y-strain (TcII). **Z3**: *T. cruzi* zymodeme 3, corresponding to DTUs TcIII and TcIV; CanIII. **M**: *T. cruzi/T. rangeli* mixed-infection; sample TBR1413. **NC**: Negative control (ddH_2_0 as template). **L**: molecular weight ladder.

### Next Generation Sequencing

Sequencing was performed on an Illumina MiSeq instrument using a 2 × 300 bp (Reagent Kit version 3) following a custom protocol [61]. Amplicon datasets were processed with Sickle for trimming (-q 20) [62]. Trimmed sequences were denoised with BayesHammer [63] and overlapping of forward and reverse sequences was performed using Pandaseq [64]. Duplicated sequences, singletons and chimeras were removed in Usearch7 and with ChimeraSlayer [65]. Haplotype cluster consensus sequences were generated *de novo* in Usearch [66]. Parasite species and DTUs were assigned to each haplotype based on mini-exon sequences available from GenBank at NCBI (S1 and S2 Tables for cloned reference strains and Ecuadorian samples, respectively).

### Richness and diversity analysis

Richness was defined as the total number of haplotypes found in each sample. As mentioned previously, samples with mixed infections showed a two-band pattern in agarose gels. For these samples, gel excision rendered two sequencing templates and the information obtained was merged (addition of reads from each template) prior to calculating diversity. Shannon Index was defined as -Σ(Piln[Pi]) [67].

### Statistical analysis

An analysis of variance (ANOVA) followed by Tukey’*s post-hoc* test was performed when data showed normality and homoscedasticity. When data violated normality, a Kruskal-Wallis test was employed. A p-value of ≤0.05 was considered significant.

## Results

### Concordance between mini-exon sequence diversity and DTU identity

The monoclonal reference strains included in the study were Chilec22 (TcI), M5521 (TcIII), Arma18 (TcIII), Saimiri13 (TcIV), Para6 (TcV), Para7 (TcVI), Chaco9 (TcVI) and LHVA (TcVI) [40, 51, 68]. The mini-exon gene sequence was analyzed for each of these monoclonal reference strains, revealing a predominant haplotype (~98% of sequences) with one or few different sequences in most samples. We note, however, that many haplotypes were shared at low abundance among multiple DTUs (S1 Table and Fig 4). High abundance (>100 reads) discriminatory sequence types identified from reference strains were used to impute DTU identities among Ecuadorian samples.

**Fig 4.**
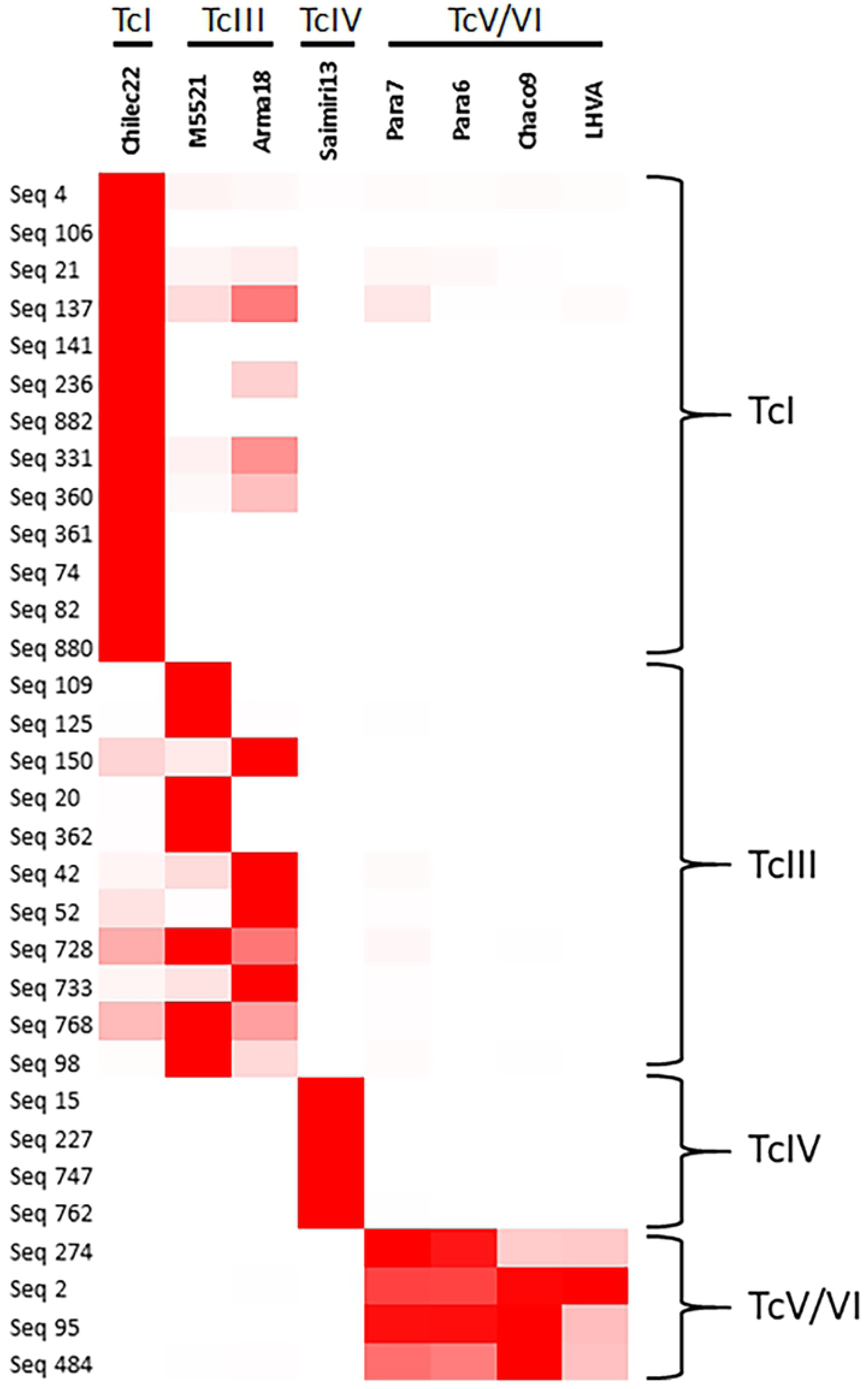
Distinctive sequence types derived from monoclonal samples. Reference strains are displayed on the x-axis. The y-axis shows the distinctive sequence types identified in this study and their corresponding DTU assignment based on similarity with GenBank reference sequences (>98%). Color intensity (red highest, white lowest) indicates the number of reads for each sequence type in different strains. The 32 sequence types displayed in the figure are those more distinctively associated with a given DTU, selected from 151 sequence types identified; selected sequences showed ≥195 reads in at least one sample.

### Genotyping by multiplex PCR and NGS

Among the 46 samples analyzed by multiplex PCR, agarose electrophoresis patterns indicated that 39.1% (18) of the samples were co-infected with *T. cruzi* and *T. rangeli*, while 54.3% (25) presented infection exclusively by *T. cruzi* and 6,5% (3) were infected only by *T. rangeli*. Meanwhile, NGS was performed only in 42 samples from the Ecuadorian dataset; four samples were not sequenced due to technical issues. An additional nine samples were excluded from further analysis because they did not reach the 2000 reads threshold, indicating insufficient depth. Among the 33 samples for which NGS was successfully performed, taxonomic assignment in GenBank identified sequence types associated with both *T. cruzi* and *T. rangeli* in 23 samples, while only 9 samples were deemed to contain exclusively *T. cruzi*-like sequences and only one sample exclusively *T. rangeli* sequences. Remarkably, sequence types matching DTU TcIV were found in samples TBR1410, TBR1422, TBR1445, TBR1475, TBR1492 and TBR1510 (Table 1 and Fig 5).

**Table 1.**
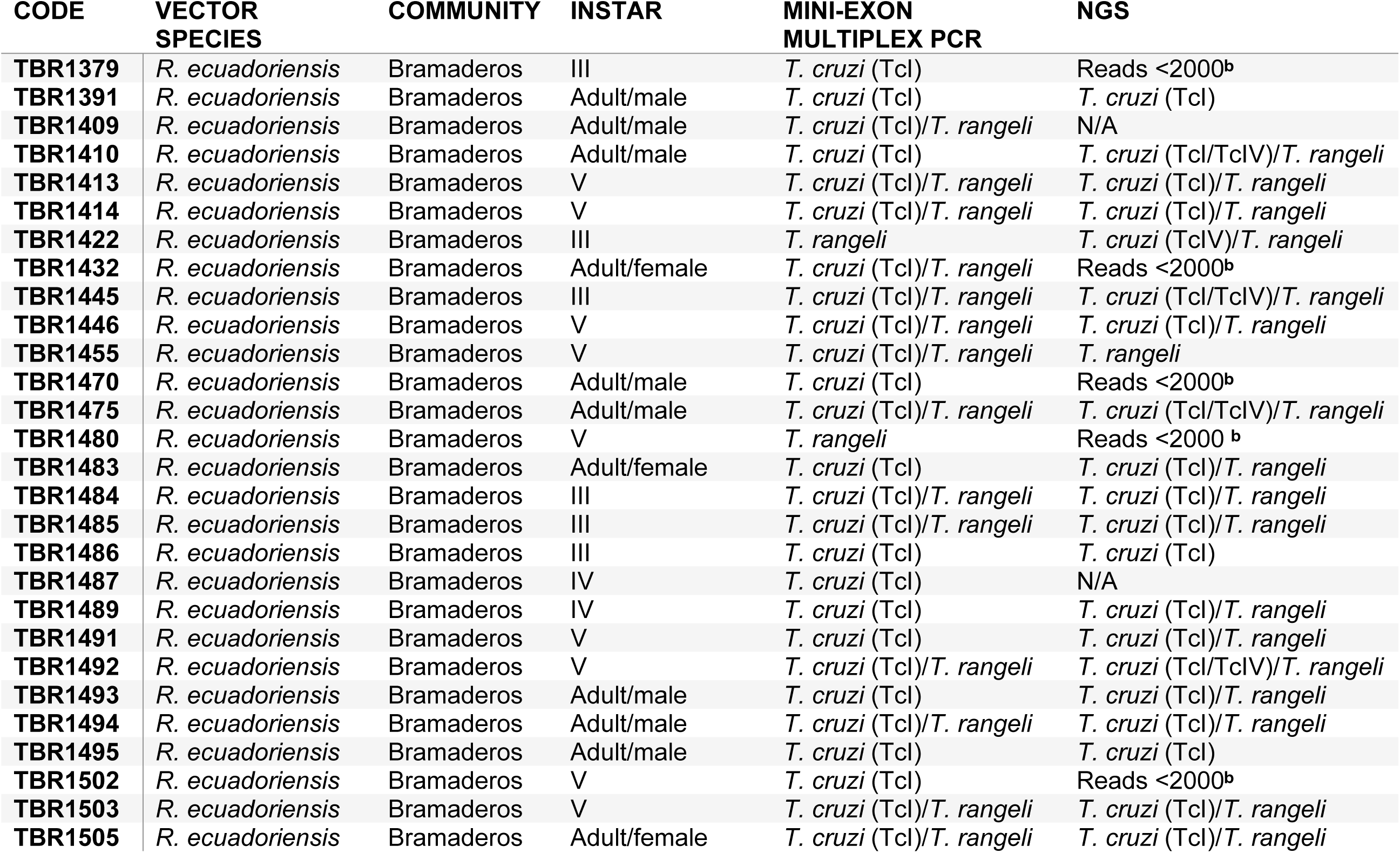
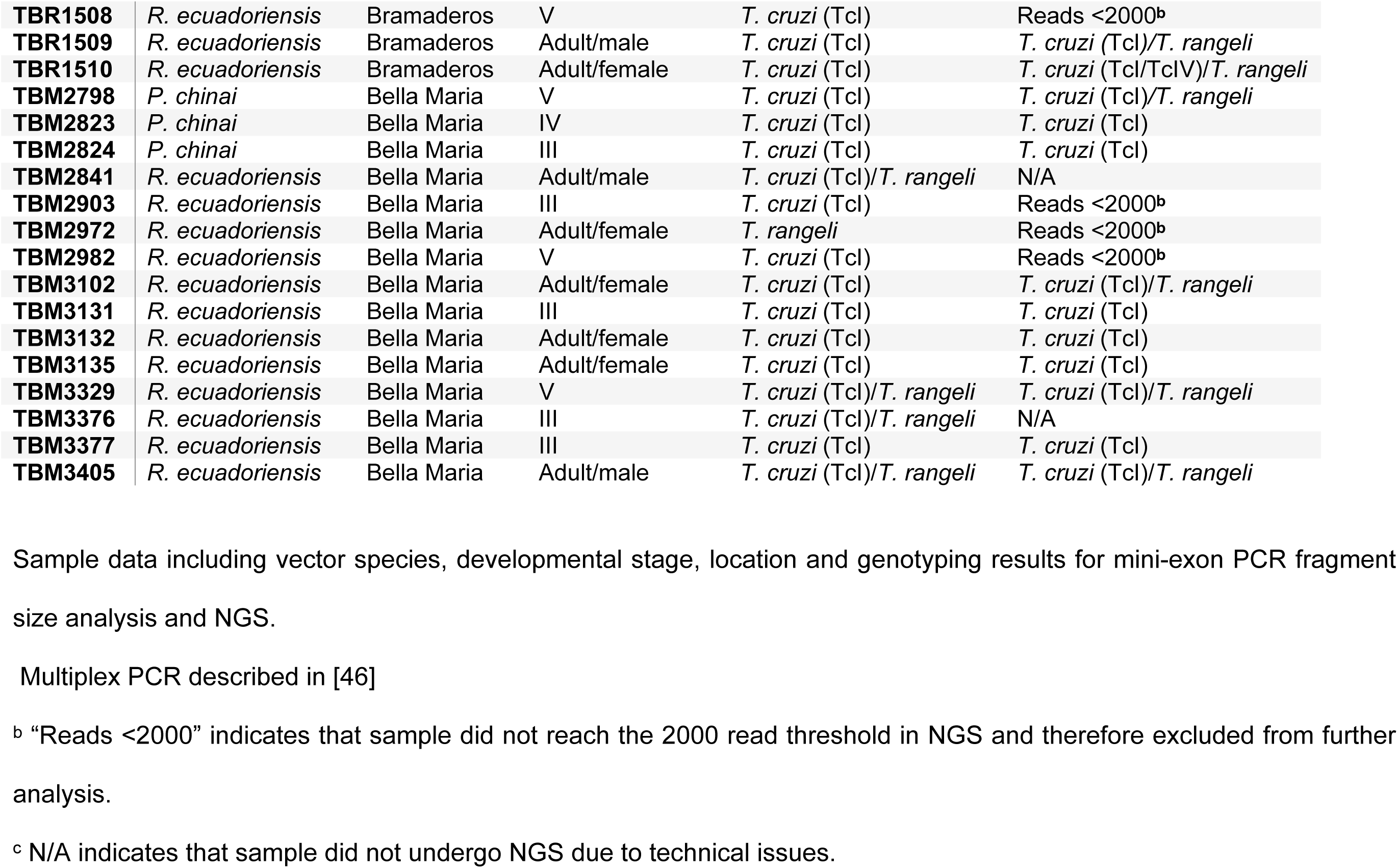
Samples employed in the study and genotyping results.

| CODE | VECTOR<br>SPECIES | COMMUNITY | INSTAR | MINI-EXON<br>MULTIPLEX PCR | NGS |
| --- | --- | --- | --- | --- | --- |
| TBR1379 | *R. ecuadoriensis* | Bramaderos | III | *T. cruzi* (Tcl) | Reads <2000<sup>b</sup> |
| TBR1391 | *R. ecuadoriensis* | Bramaderos | Adult/male | *T. cruzi* (Tcl) | *T. cruzi* (Tcl) |
| TBR1409 | *R. ecuadoriensis* | Bramaderos | Adult/male | *T. cruzi* (Tcl)/*T. rangeli* | N/A |
| TBR1410 | *R. ecuadoriensis* | Bramaderos | Adult/male | *T. cruzi* (Tcl) | *T. cruzi* (Tcl/TclV)/*T. rangeli* |
| TBR1413 | *R. ecuadoriensis* | Bramaderos | V | *T. cruzi* (Tcl)/*T. rangeli* | *T. cruzi* (Tcl)/*T. rangeli* |
| TBR1414 | *R. ecuadoriensis* | Bramaderos | V | *T. cruzi* (Tcl)/*T. rangeli* | *T. cruzi* (Tcl)/*T. rangeli* |
| TBR1422 | *R. ecuadoriensis* | Bramaderos | III | *T. rangeli* | *T. cruzi* (TclV)/*T. rangeli* |
| TBR1432 | *R. ecuadoriensis* | Bramaderos | Adult/female | *T. cruzi* (Tcl)/*T. rangeli* | Reads <2000<sup>b</sup> |
| TBR1445 | *R. ecuadoriensis* | Bramaderos | III | *T. cruzi* (Tcl)/*T. rangeli* | *T. cruzi* (Tcl/TclV)/*T. rangeli* |
| TBR1446 | *R. ecuadoriensis* | Bramaderos | V | *T. cruzi* (Tcl)/*T. rangeli* | *T. cruzi* (Tcl)/*T. rangeli* |
| TBR1455 | *R. ecuadoriensis* | Bramaderos | V | *T. cruzi* (Tcl)/*T. rangeli* | *T. rangeli* |
| TBR1470 | *R. ecuadoriensis* | Bramaderos | Adult/male | *T. cruzi* (Tcl) | Reads <2000<sup>b</sup> |
| TBR1475 | *R. ecuadoriensis* | Bramaderos | Adult/male | *T. cruzi* (Tcl)/*T. rangeli* | *T. cruzi* (Tcl/TclV)/*T. rangeli* |
| TBR1480 | *R. ecuadoriensis* | Bramaderos | V | *T. rangeli* | Reads <2000<sup>b</sup> |
| TBR1483 | *R. ecuadoriensis* | Bramaderos | Adult/female | *T. cruzi* (Tcl) | *T. cruzi* (Tcl)/*T. rangeli* |
| TBR1484 | *R. ecuadoriensis* | Bramaderos | III | *T. cruzi* (Tcl)/*T. rangeli* | *T. cruzi* (Tcl)/*T. rangeli* |
| TBR1485 | *R. ecuadoriensis* | Bramaderos | III | *T. cruzi* (Tcl)/*T. rangeli* | *T. cruzi* (Tcl)/*T. rangeli* |
| TBR1486 | *R. ecuadoriensis* | Bramaderos | III | *T. cruzi* (Tcl) | *T. cruzi* (Tcl) |
| TBR1487 | *R. ecuadoriensis* | Bramaderos | IV | *T. cruzi* (Tcl) | N/A |
| TBR1489 | *R. ecuadoriensis* | Bramaderos | IV | *T. cruzi* (Tcl) | *T. cruzi* (Tcl)/*T. rangeli* |
| TBR1491 | *R. ecuadoriensis* | Bramaderos | V | *T. cruzi* (Tcl) | *T. cruzi* (Tcl)/*T. rangeli* |
| TBR1492 | *R. ecuadoriensis* | Bramaderos | V | *T. cruzi* (Tcl)/*T. rangeli* | *T. cruzi* (Tcl/TclV)/*T. rangeli* |
| TBR1493 | *R. ecuadoriensis* | Bramaderos | Adult/male | *T. cruzi* (Tcl) | *T. cruzi* (Tcl)/*T. rangeli* |
| TBR1494 | *R. ecuadoriensis* | Bramaderos | Adult/male | *T. cruzi* (Tcl)/*T. rangeli* | *T. cruzi* (Tcl)/*T. rangeli* |
| TBR1495 | *R. ecuadoriensis* | Bramaderos | Adult/male | *T. cruzi* (Tcl) | *T. cruzi* (Tcl) |
| TBR1502 | *R. ecuadoriensis* | Bramaderos | V | *T. cruzi* (Tcl) | Reads <2000<sup>b</sup> |
| TBR1503 | *R. ecuadoriensis* | Bramaderos | V | *T. cruzi* (Tcl)/*T. rangeli* | *T. cruzi* (Tcl)/*T. rangeli* |
| TBR1505 | *R. ecuadoriensis* | Bramaderos | Adult/female | *T. cruzi* (Tcl)/*T. rangeli* | *T. cruzi* (Tcl)/*T. rangeli* |
| **TBR1508** | *R. ecuadoriensis* | Bramaderos | V | *T. cruzi* (Tcl) | Reads <2000<sup>b</sup> |
| **TBR1509** | *R. ecuadoriensis* | Bramaderos | Adult/male | *T. cruzi* (Tcl) | *T. cruzi* (Tcl)/*T. rangeli* |
| **TBR1510** | *R. ecuadoriensis* | Bramaderos | Adult/female | *T. cruzi* (Tcl) | *T. cruzi* (Tcl/TclV)/*T. rangeli* |
| **TBM2798** | *P. chinai* | Bella Maria | V | *T. cruzi* (Tcl) | *T. cruzi* (Tcl)/*T. rangeli* |
| **TBM2823** | *P. chinai* | Bella Maria | IV | *T. cruzi* (Tcl) | *T. cruzi* (Tcl) |
| **TBM2824** | *P. chinai* | Bella Maria | III | *T. cruzi* (Tcl) | *T. cruzi* (Tcl) |
| **TBM2841** | *R. ecuadoriensis* | Bella Maria | Adult/male | *T. cruzi* (Tcl)/*T. rangeli* | N/A |
| **TBM2903** | *R. ecuadoriensis* | Bella Maria | III | *T. cruzi* (Tcl) | Reads <2000<sup>b</sup> |
| **TBM2972** | *R. ecuadoriensis* | Bella Maria | Adult/female | *T. rangeli* | Reads <2000<sup>b</sup> |
| **TBM2982** | *R. ecuadoriensis* | Bella Maria | V | *T. cruzi* (Tcl) | Reads <2000<sup>b</sup> |
| **TBM3102** | *R. ecuadoriensis* | Bella Maria | Adult/female | *T. cruzi* (Tcl) | *T. cruzi* (Tcl)/*T. rangeli* |
| **TBM3131** | *R. ecuadoriensis* | Bella Maria | III | *T. cruzi* (Tcl) | *T. cruzi* (Tcl) |
| **TBM3132** | *R. ecuadoriensis* | Bella Maria | Adult/female | *T. cruzi* (Tcl) | *T. cruzi* (Tcl) |
| **TBM3135** | *R. ecuadoriensis* | Bella Maria | Adult/female | *T. cruzi* (Tcl) | *T. cruzi* (Tcl) |
| **TBM3329** | *R. ecuadoriensis* | Bella Maria | V | *T. cruzi* (Tcl)/*T. rangeli* | *T. cruzi* (Tcl)/*T. rangeli* |
| **TBM3376** | *R. ecuadoriensis* | Bella Maria | III | *T. cruzi* (Tcl)/*T. rangeli* | N/A |
| **TBM3377** | *R. ecuadoriensis* | Bella Maria | III | *T. cruzi* (Tcl) | *T. cruzi* (Tcl) |
| **TBM3405** | *R. ecuadoriensis* | Bella Maria | Adult/male | *T. cruzi* (Tcl)/*T. rangeli* | *T. cruzi* (Tcl)/*T. rangeli* |

Sample data including vector species, developmental stage, location and genotyping results for mini-exon PCR fragment size analysis and NGS.
Multiplex PCR described in [46]
<sup>b</sup> “Reads <2000” indicates that sample did not reach the 2000 read threshold in NGS and therefore excluded from further analysis.
<sup>c</sup> N/A indicates that sample did not undergo NGS due to technical issues.

**Fig 5.**
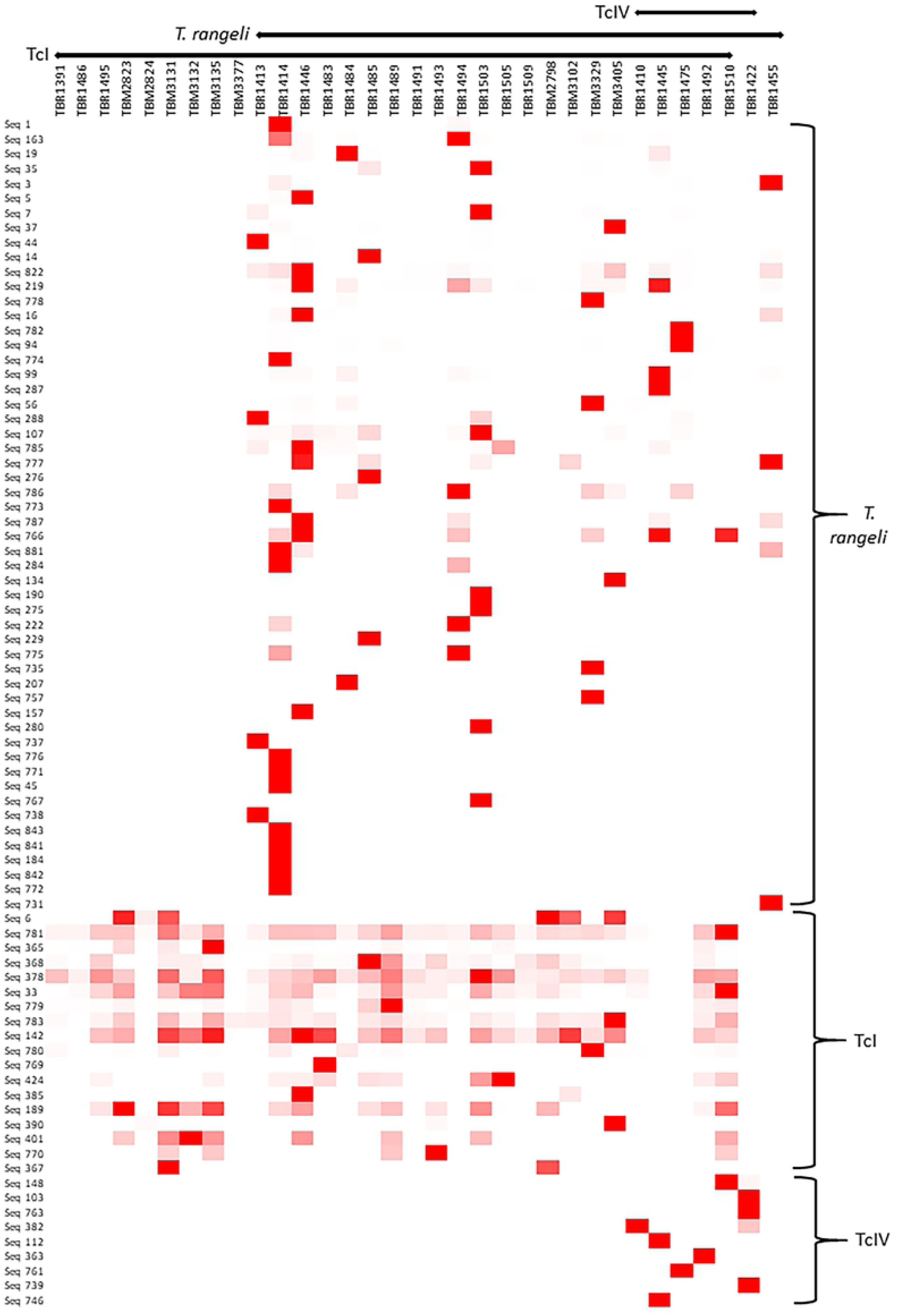
Sequence types associated with *T. cruzi* and/or *T. rangeli* amplified from intestinal content of Ecuadorian triatomines. The y-axis shows the sequence types and their assigned DTU, according to their similarity with GenBank reference sequences (>98%). Samples are shown in the x-axis, indicating their assigned species and DTU. Color intensity (red highest, white lowest) indicates the number of reads for each sequence type in each sample. Six sequences were excluded from the figure do to ≤125 reads in at least one sample (Seq 361, Seq 862 and Seq 404) or ubiquitous throughout all samples and less informative.

Sequence types distinctively associated with monoclonal strains TcI, TcIII, TcIV and TcV/VI (Fig 4), as well with selected sequence types derived from Ecuadorian strains, were used for the construction of a Neighbor-Joining tree. The tree topology shows three robustly supported clusters, with TcI-like sequences in one branch, separated from TcIV-like and TcV/VI-like sequences (Fig 6).

**Fig 6.**
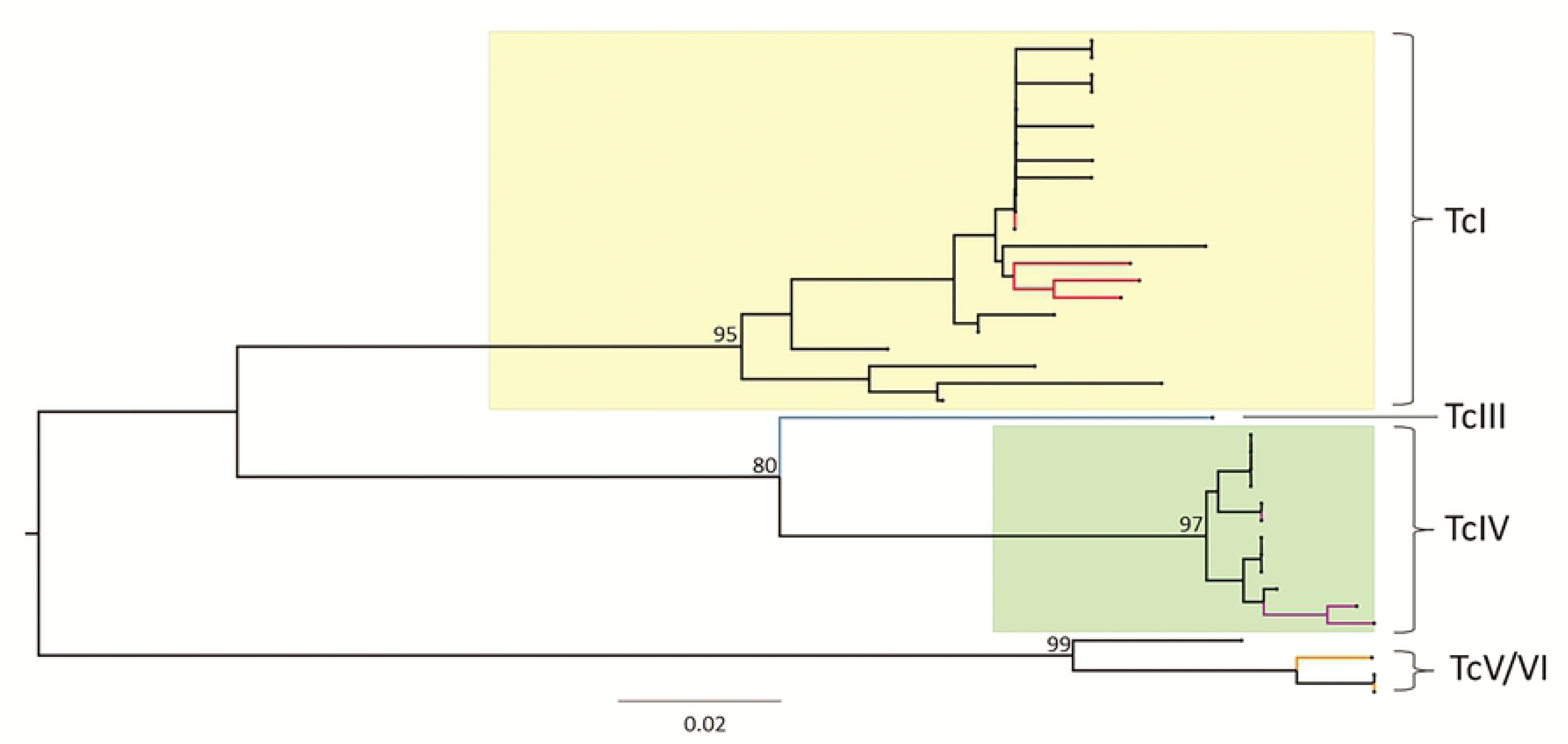
Neighbor-joining analysis of selected mini-exon sequence types derived from Ecuadorian samples and monoclonal reference strains. Midpoint-rooted tree generated in MEGA7. Alignment performed with ClustalW. Positions from nucleotide 99 to 355 were excluded due to indels and poor alignment. Three clusters are supported by 1000 replicates for Bootstrap values of 95% for TcI, 97% for TcIV and 99% for TcV/VI. Colored branches represent monoclonal reference strains sequences (red for TcI, blue for TcIII, violet for TcIV and orange for TcV/VI).

### Parasite richness and diversity

Chao richness and Shannon diversity indexes were calculated for each parasite species within each sample. No significant differences in richness or diversity were found between samples from the two studied localities (Bramaderos and Bella Maria) for either parasite species. In terms of diversity, significant differences were detected for *T. cruzi* between instar V nymphs and adult triatomines [F(2,26) = 5.329, p = 0.011] (Fig 7). No other significant differences were detected when comparing diversity/richness indexes corresponding to both parasite species across vector developmental stages, including data for males and females analyzed independently, or combined as a merged group identified as adults.

**Fig 7.**
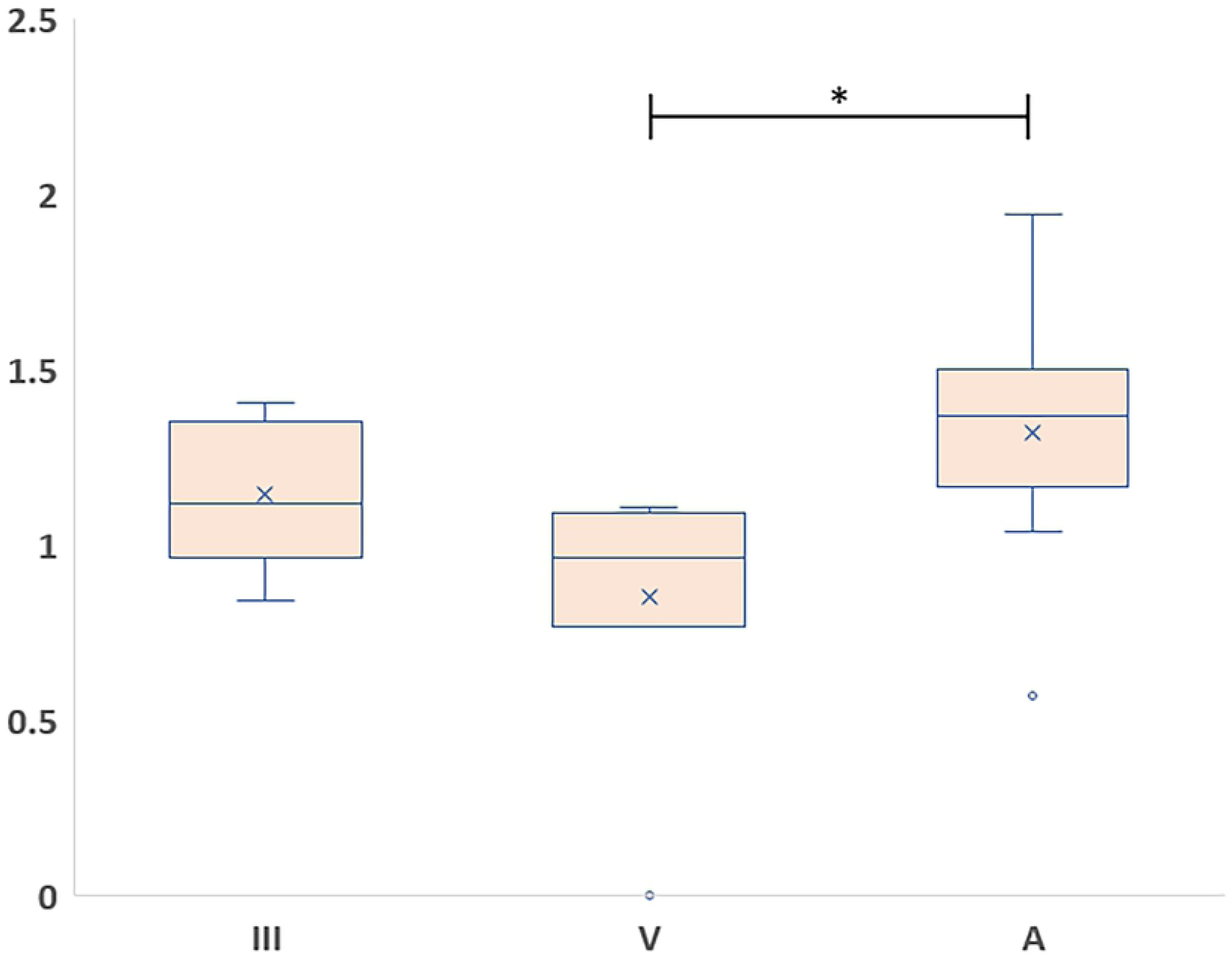
Diversity of *T. cruzi* sequence haplotypes of the non-transcribed spacer region of the mini-exon gene across different developmental stages of the vector. The Shannon diversity Index calculated for *T. cruzi* haplotypes for both localities is shown in the y-axis. Vector developmental stages shown in the x-axis (**III** and **V**: third and fifth instar, respectively. **A**: adults, males and females combined). * indicates *p* < 0.05. Data corresponding to samples TBM2823 and TBR1489 were excluded from this analysis because they are the only representatives of instar IV. TBR1446 and TBR1503 were also excluded to avoid bias because the sequence reads from these samples has been merged across four sequencing reactions.

## Discussion

We analyzed 46 intestinal content DNA samples from triatomine bugs collected in two rural communities of Loja province, southern Ecuador. Samples were analyzed via three different molecular methodologies. All samples selected for the study had been previously characterized as infected with *T. cruzi* based on PCR product size polymorphism of kinetoplast minicircle sequences, with no evidence of co-infection with the sympatric sister species *T. rangeli*. The sample set was subsequently analyzed by two additional molecular techniques – first by PCR product size polymorphism analysis of the non-transcribed spacer region of the mini-exon gene, then by deep sequencing of these mini-exon amplicons. Incongruences were detected among the results obtained by the three methods, and our results suggest that simple amplicon visualization of kinetoplast minicircle sequences may not suffice for detection of *T. cruzi-T. rangeli* co-infections.

The PCR-based method for mini-exon gene analysis without sequencing [49] uncovered 18 instances of *T. cruzi/T. rangeli* co-infections in our sample panel which the kinetoplast minicircle PCR [60] did not identify. Mixed infections of *T. cruzi* and *T. rangeli* in vector and mammalian hosts are frequent [6–8, 69] and several groups have reported that minicircle kinetoplast PCR product visualization techniques are inadequate to identify them [69–72]. Previous studies have applied this same method in southern Ecuador and reported ~10% prevalence of *T. rangeli* infection in sampled triatomines and mammals and just ~1.25% in co-infection with *T. cruzi* [12]. In stark contrast, with sequencing *T. rangeli*-like sequences were detected in 23 out of 33 (~70%) infected samples characterized in our study. Our sample panel was heavily biased toward *R. ecuadoriensis* (43 out of 46 DNA samples were isolated from this vector species), one of the two most important vectors of CD in the study region and in Ecuador at large [20]. A close association between *T. rangeli* and *Rhodnius* has been established in the past, where 12 out of 15 species of the genus have shown vectorial capacity [73]. Our results suggest that previous reports regarding *T. rangeli* infection in *R. ecuadoriensis* may underestimate the presence of this parasite in southern Ecuador, where a wider distribution of *T. rangeli* in Loja, and perhaps the rest of the country, may have been overlooked.

Sequencing of the non-transcribed spacer of the mini-exon gene has proven useful in phylogenetic analysis and has provided valuable insights for DTU discrimination while analyzing *T. cruzi* intra-strain variability [40, 50, 53, 54, 74]. Recently, deep sequencing of the non-transcribed spacer has revealed high levels of *T. cruzi* diversity in the intestinal content of *T. dimidiata* [31]. Similarly, Herrera and colleagues recently employed NGS sequencing of the mini-exon gene to analyze the genetic diversity of *T. cruzi* naturally infecting a group of captive non-human primates in the southern USA [51]. In our study, 12 *T. cruzi* haplotypes per triatomine were detected on average. However, we do note from our analysis of monoclonal reference samples that mini-exon sequences may not accurate resolve *T. cruzi* phylogenetic diversity – reflecting to an extent the lack of reliability of this locus in identifying DTUs. From monoclonal strains haplotypes our data shows that certain low abundance sequence types are shared by almost all DTUs – with important implications for those using the mini-exon splice leader locus to define intra-host DTU diversity [31]. It is therefore difficult to say with confidence that the high levels of *T. cruzi* diversity found in our samples result from multiclonal infections.

TcI is the predominant DTU in Ecuador [7, 19, 41, 42]. As expected, TcI-like haplotypes were widely represented in the deep sequencing analysis of the mini-exon gene performed for the Ecuadorian samples. Sequences most abundant in the DTU TcIV clone were amplified from six of the Ecuadorian samples. If this does indeed represent the presence of this DTU in our samples, this constitutes the first report of TcIV in Loja Province and is testament to the power of amplicon sequencing to uncover ‘hidden’ *T. cruzi* diversity, especially in TBR1472 where this TcIV clone occurs as a mixed infection alongside sequences more closely related to TcI. Neighbor-joining analysis clustered the Ecuadorian TcIV haplotypes with discriminatory TcIV sequences available from monoclonal strains with known DTU identity. Strong bootstrap values support this finding. Only two previous reports of DTUs different from TcI in Ecuador exist. One of them, from the early 2000s, employed multilocus enzyme electrophoresis to identify zymodemes II and III [43]. The second report came from serological evidence using a TcII/V/VI-specific epitope [44]. Our findings expand the distribution of DTU TcIV to the South of Ecuador, where its presence had not previously been reported.

No major differences were found among the studied localities in terms of parasite genetic richness or diversity. Both communities are only 28.3 km apart and infected livestock, small mammals, passive transportation of triatomines or human movement may cause active parasite dispersal, homogenizing the parasite populations.

Our data suggest a tendency where several clones of *T. cruzi* accumulate in late developmental stages of the vector. Based on the estimation of infection events, Dumonteil and colleagues found similar results in the intestinal content of *T. dimidiata* [31]. Altogether, this evidence could support a direct association between parasite diversity and the time vectors are exposed to infection sources. It is noteworthy that this effect has been noted in the invertebrate vector (*T. dimidiata, R. ecuadoriensis*), while evidence of its occurrence in mammalian hosts has yet to be established [29, 30].

Our results also indicate that techniques based solely on PCR amplification of minicircle or mini-exon sequences are not optimal for detection and characterization of mixed infections of kinetoplastid parasites in triatomines. Employing deep sequencing of the non-transcribed spacer region of the mini-exon gene, we have encountered ~70% of studied triatomines to be co-infected with *T. cruzi* and *T. rangeli*, a much higher prevalence for *T. rangeli* infection than previously reported for vectors in the study region. Additionally, we provide the first possible report for the presence of DTU TcIV in southern Ecuador.

In conclusion, we demonstrate the power of NGS to scrutinize parasite populations with high resolution. However, we do note that the mini-exon marker must be approached with caution, and further deep sequencing analyses targeted at additional markers – potentially conserved house-keeping genes [75] – are required for confirmation. Nonetheless, the information presented in this report is of value in the characterization of the dynamics of *T. cruzi* transmission in southern Ecuador and must be considered in disease control efforts in the region.

## Acknowledgements

Special thanks to Anita Villacís from CISeAL’s Medical Entomology unit; to Alejandra Zurita and Sofía Ocaña for parasite and DNA isolation, information regarding infection status based on kinetoplast PCR gel electrophoresis, and associated metadata; and to César Yumiseva for assistance in the generation of maps. Special thanks to Julie Galbraith at Glasgow Polyomics for assistance with Next Generation Sequencing, and to Camila Cilveti, Christopher Garcia and Andrea Vela from CISeAL.

## Supporting information

**S1 Table. DTU assignment for monoclonal reference strain sequences based on similarity with NCBI entries**

**S2 Table. Species and DTU assignment for Ecuadorian sample sequences based on similarity with NCBI entries and with DTU distinctive sequences retrieved from monoclonal strains**

